# Multifunctional nanozyme therapy accelerates hematoma clearance and attenuates genome damage and senescence after intracerebral hemorrhage

**DOI:** 10.64898/2026.03.12.711396

**Authors:** Vikas H. Malojirao, Andrei M. Mikheev, Manohar Kodavati, Anton V. Liopo, Paul J. Derry, Elias Perli, Karthik Mouli, Shu Zhang, Zhonglin Liu, Xiufeng Tang, Gavin W. Britz, Philip J. Horner, Thomas A. Kent, Muralidhar L. Hegde

## Abstract

Intracerebral hemorrhage (ICH) is a devastating form of stroke characterized by rapid hematoma formation in the brain, resulting in multiple pathological events due to mass effect and toxicity of extravasated blood and its blood products. ICH leads to poor long-term outcomes despite advances in hematoma management, largely due to secondary injury mechanisms. Hemin and iron released in the peri-hematomal environment trigger genome damage, transient senescence, and inflammatory signaling that may initially limit ferroptosis but ultimately contribute to persistent neurodegeneration. Given the multiple pathological events initiated following ICH, it is not surprising that no single neuroprotective strategy has been effective. In this study, we investigated these interconnected pathways in a rodent model of ICH and evaluated the therapeutic potential of DEF-OAC-PEG, a pleiotropic synthetic oxidized carbon nano-enzyme that has catalytic mitochondrial and cellular protective actions, covalently bonded to the iron chelator deferoxamine and shown in our previous work to have strong *in vitro* protective effects against hemin and iron toxicity and *in vivo* evidence of reduction in genome damage. Here, we examined mechanisms of action in an *in vivo* ICH mouse model. Autologous whole blood injection into the mouse brain striatum induced robust astroglial and microglial activation, increased neuronal Heme Ooxygenase-1 expression, and DNA damage and senescence in neurons and oligodendrocytes. Systemic intraperitoneal administration of DEF-OAC-PEG, initiated 3 hours after ICH, resulted in robust brain penetration in wild-type mice, with preferential accumulation in peri-hematomal regions of ICH mice. Surprisingly, nanozyme treatment produced a rapid, significant acceleration of hematoma clearance compared with untreated ICH animals. This effect was associated with enhanced detection of CD68-positive microglia/macrophages, which also showed internalized nanozymes, suggesting that nanozyme promotes immune-mediated hematoma resolution. Importantly, DEF-OAC-PEG also markedly attenuated ICH-induced DNA damage and senescence in neurons and oligodendrocytes. Together, these findings identify genome instability and senescence as key consequences of hemorrhagic brain injury and demonstrate that multifunctional nanozyme therapy can simultaneously promote hematoma resolution and mitigate secondary neurodegenerative injury following ICH.

## Introduction

Intracerebral hemorrhage (ICH) is one of the most devastating forms of stroke, accounting for approximately 25-30% of all stroke cases but contributing disproportionately to mortality and long-term neurological disability^1,2^. Despite improvements in neurosurgical management and supportive care, clinical outcomes for ICH remain poor, largely because effective therapies targeting the secondary injury processes that follow the initial hemorrhage are lacking^3,4^. Following vascular rupture, extravasated blood rapidly accumulates within the brain parenchyma, forming a hematoma that exerts mechanical compression on surrounding tissue and triggers a cascade of biochemical and cellular responses^5,6^. These secondary injury mechanisms, including neuroinflammation, mitochondrial dysfunction, and progressive neuronal damage, continue for days to weeks after the initial insult and represent critical therapeutic targets^7-9^.

Among the toxic mediators generated during hematoma degradation, hemin and iron derived from hemoglobin breakdown play central roles in driving tissue injury in the peri-hematomal region^10^. Hemin, a protoporphyrin released from lysed erythrocytes, accumulates at high concentrations and promotes oxidative stress, lipid peroxidation, and neuronal toxicity^11^. Heme metabolism is further amplified by the induction of heme oxygenase-1 (HO-1), which degrades hemin into biliverdin, carbon monoxide, and ferrous iron^12,13^. Because HO-1 is not constitutively expressed in brain neurons, its induction post hemin exposure lags many toxic effects of hemin. Although this pathway initially serves a protective function by detoxifying free heme, the subsequent accumulation of labile iron can overwhelm cellular iron-handling systems, promoting Fenton reaction and ferroptotic cell death^14-16^. In parallel, iron can directly disrupt mitochondrial metabolism and impair redox-sensitive DNA repair pathways, amplifying neuronal vulnerability following hemorrhagic injury^17^. Despite extensive evidence implicating hemin and iron toxicity in ICH pathology, the molecular mechanisms linking these insults to long-term neuronal dysfunction remain incompletely understood^18,19^.

Recent evidence suggests that genome instability and cellular senescence may represent critical yet underappreciated components of hemorrhagic brain injury. We have demonstrated that exposure to hemin induces persistent nuclear and mitochondrial DNA damage in neurons and vascular cells, triggering a robust DNA damage response (DDR)^20,21^. These events promote the induction of a senescence-like state characterized by activation of p21-dependent pathways and the emergence of senescence-associated inflammatory signaling^21^. Interestingly, early induction of transient senescence may represent an adaptive stress response that allows cells to buffer acute heme and iron toxicity, thereby protecting against ferroptotic cell death^21^. However, prolonged or repeated exposure to hemin appears to shift this response toward persistent pathological senescence, accompanied by sustained inflammatory signaling and chronic neurodegenerative processes^21^. These findings suggest that hemorrhagic brain injury involves a complex interplay between mitochondrial dysfunction and free radical excess, iron toxicity, genome instability, and senescence-associated inflammation^21^.

Given the multifactorial nature of these injury pathways, therapeutic strategies targeting a single mechanism have shown limited success^4^. Traditional antioxidant approaches or iron chelation therapies alone have demonstrated modest benefits in experimental models but are limited by poor cellular uptake, unfavorable stoichiometry and short half-life in the face of a cascade of radicals, and inadequate targeting of intracellular injury pathways, particularly within mitochondria where the early oxidative radicals and damage occurs^22-26^. Therefore, therapeutic platforms capable of simultaneously addressing multiple pathological mechanisms may provide a more effective strategy for mitigating secondary injury after ICH.

To address these challenges, we previously developed a multifunctional (pleiotropic) carbon-based nanozyme capable of simultaneously preventing hemin-induced cellular senescence and iron-mediated ferroptosis in cultured neurons^20^. This nanozyme is based on PEGylated oxidized activated charcoal (PEG-OAC), a catalytically active nanomaterial with intrinsic redox-modulating properties that mimics superoxide dismutase enzyme activity and promotes cellular metabolic flexibility by facilitating the oxidation of NADH to NAD+ and resilience. It is avidly taken up by multiple cell types and co-localizes with mitochondria and other cellular membranes. In the presence of hemin (and consequently its liberated iron), PEG-OAC effectively reduced senescence, but unleashed ferroptosis ^20,21^. To enhance protection against iron toxicity, we conjugated the clinically used iron chelator deferoxamine (DEF) to the PEG-OAC scaffold to generate the multifunctional nanozyme DEF-OAC-PEG^20,21,27^. This design integrates the nanozyme’s activity with iron chelation while improving cellular uptake and mitochondrial enrichment afforded by the PEG-OAC core^28^. In vitro studies demonstrated that DEF-OAC-PEG reduces radical damage, preserves nuclear and mitochondrial DNA integrity, and prevents lipid peroxidation associated with ferroptosis following hemin exposure^20,21^.

In the present study, we extend these findings to an in vivo setting by investigating the pathological interplay between genome damage, cellular senescence, and iron-mediated toxicity in a rodent model of ICH and evaluating the therapeutic potential of DEF-OAC-PEG nanozyme treatment. Using a stereotactic autologous whole blood injection model of ICH, we demonstrate that hemorrhagic injury induces robust glial activation, neuronal HO-1 expression, and pronounced DNA damage and senescence in neurons and oligodendrocytes. Systemically administered DEF-OAC-PEG penetrates the brain, localizes to mitochondria, and is internalized by phagocytic microglia/macrophages in peri-hematomal regions. Remarkably, nanozyme treatment significantly reduced hematoma volume and accelerated hematoma clearance, an effect associated with enhanced accumulation of phagocytic CD68-positive cells. In parallel, nanozyme therapy attenuated DNA damage and senescence in neural cell populations. Together, these findings indicate that genome instability and senescence are key consequences of hemorrhagic brain injury and suggest that multifunctional nanozyme strategies may represent a promising therapeutic approach for promoting hematoma resolution and mitigating secondary neurodegenerative injury following ICH.

## Materials and Methods

### Reagents and Experimental Materials

DEF-OAC-PEG nanozymes (deferoxamine-conjugated PEGylated oxidized activated charcoal nanoparticles) were used for therapeutic administration in the experimental intracerebral hemorrhage (ICH) model. Isoflurane (Fluriso, NDC 13985-528-60) was used for anesthesia during surgical procedures. Phosphate-buffered saline (PBS; Gibco, Thermo Fisher Scientific, Cat# 10010-023) and 4% paraformaldehyde (PFA; Sigma-Aldrich, Cat# P6148) were used for tissue perfusion and fixation. Brain tissues were cryoprotected using sucrose solution (Sigma-Aldrich, Cat# S0389) prior to freezing and cryosectioning. Triton X-100 (Sigma-Aldrich, Cat# T8787) was used for tissue permeabilization, and bovine serum albumin (BSA; Sigma-Aldrich, Cat# A9647) was used for blocking during immunostaining procedures. Primary antibodies used in this study included anti-MAP2 (neuronal marker; Abcam, Cat# ab5392), anti-Olig2 (oligodendrocyte marker; Millipore, Cat# AB9610), anti-GFAP (astrocyte marker; Abcam, Cat# ab4674), anti-Iba1 (microglial marker; Abcam, Cat# ab178846), anti-CD68 (phagocytic microglia/macrophage marker; Bio-Rad, Cat# MCA1957), anti-NeuN (neuronal marker; Abcam, Cat# ab177487), anti-γH2AX (DNA double-strand break marker; Cell Signaling Technology, Cat# 9718), and anti-HO-1 (heme oxygenase-1; Abcam, Cat# ab13248). Fluorophore-conjugated secondary antibodies used for immunofluorescence included Alexa Fluor 488 goat anti-mouse IgG (Thermo Fisher Scientific, Cat# A-11001), Alexa Fluor 488 goat anti-rabbit IgG (Thermo Fisher Scientific, Cat# A-11008), Alexa Fluor 647 goat anti-mouse IgG (Thermo Fisher Scientific, Cat# A-21235), Alexa Fluor 647 goat anti-rabbit IgG (Thermo Fisher Scientific, Cat# A-21245), and Alexa Fluor 647 goat anti-chicken IgG (Thermo Fisher Scientific, Cat# A-21449). For immunohistochemistry, biotinylated secondary antibodies were used together with an avidin–biotin complex (ABC) detection system, followed by visualization with 3,3′ -diaminobenzidine (DAB) substrate (Vector Laboratories, Cat# SK-4100). Nuclei were counterstained with DAPI (Invitrogen, Cat# D1306). Cellular senescence was assessed using a Senescence β -Galactosidase Staining Kit (Cell Signaling Technology, Cat# 9860), which utilizes X-gal substrate for detection of senescence-associated β-galactosidase (SA-β-Gal) activity. Stereotactic intracerebral injections were performed using a stereotactic frame (Kopf Instruments) and a 26-gauge Hamilton microsyringe (Hamilton Company), and burr holes were generated using a microsurgical drill. Coronal brain sections for gross anatomical analysis were prepared using a mouse brain matrix (Zivic Instruments, Cat# BSMAS001). Brain tissues were sectioned using a Leica CM1950 cryostat (Leica Microsystems) for histological and immunofluorescence analyses. Fluorescence and brightfield images were acquired using an Olympus slide scanner (Olympus, Tokyo, Japan) for whole-slide imaging, while high-resolution fluorescence images were obtained using a FluoView FV3000 confocal laser scanning microscope (Olympus) equipped with appropriate laser lines and objectives. In vivo brain imaging was performed using a 7T small-animal magnetic resonance imaging (MRI) system (MR Solutions, Guildford, UK), acquiring T2-weighted fast spin-echo (T2w FSE) images for hematoma visualization and lesion volume assessment. Lesion volumes were quantified with MicroDicom, and statistical analyses were performed in GraphPad Prism.

### Animals and an intracerebral hemorrhage (ICH) model

Adult C57BL/6 mice of either sex (8–12 weeks old) were used for all experiments. All procedures were approved by the Institutional Animal Care and Use Committee (IACUC) of Houston Methodist Research Institute. Mice were housed in a pathogen-free facility under controlled temperature and humidity with a 12-h light/dark cycle and provided ad libitum access to food and water. ICH was induced using a stereotactic autologous whole-blood injection method^29^. Adult mice were anesthetized with isoflurane and secured in a stereotactic frame. Following a midline scalp incision, the skull was exposed and a burr hole was drilled at stereotactic coordinates relative to bregma (0.2 mm anterior and 2.0 mm lateral). A total of 30 µL of autologous arterial blood was freshly collected and loaded into a Hamilton microsyringe. The needle was lowered vertically into the right striatum to a depth of 3.0 mm from the dural surface. An initial volume of 5 µL of whole blood was injected at a controlled rate of 2 µL/min to minimize tissue disruption. After completion of the initial injection, the needle was advanced an additional 0.7 mm and maintained in position for 5 min to facilitate clot formation and prevent reflux along the needle tract. The remaining 25 µL of blood was then injected at the same rate, followed by an additional 10 min dwell period. The needle was subsequently withdrawn slowly at a rate of 1 mm/min. The burr hole was sealed, the scalp incision was sutured, and animals were allowed to recover under temperature-controlled conditions. Sham-operated animals underwent identical surgical procedures, including stereotactic needle placement and dwell times, but without blood injection.

### Nanozyme treatment

To evaluate the effects of DEF-OAC-PEG nanozyme administration in an experimental model of ICH, eighteen C57BL/6 mice were randomly assigned to four groups (n = 6 per group): (1) sham, (2) ICH with vehicle treatment, and (3) ICH + DEF-OAC-PEG nanozymes. DEF-OAC-PEG nanozymes are deferoxamine-conjugated PEGylated oxidized activated charcoal nanoparticles engineered to combine iron-chelating activity with redox-modulating nanozyme properties and were synthesized as described in our previous studies^20,30^. Nanozymes were administered 2h post-surgery via intraperitoneal injection at a dose of 2 mg/kg body weight according to the treatment schedule illustrated in Figure 3a. At designated time points, mice were either sacrificed, and brain tissues were collected for biochemical analyses or subjected to longitudinal MRI imaging assessments. Nanozyme dose, route of administration, and timing were maintained consistently across all experimental groups.

### Brain tissue processing and sectioning

At designated time points following ICH induction, animals were deeply anesthetized and transcardially perfused with phosphate-buffered saline (PBS), followed by 4% paraformaldehyde (PFA). Brains were harvested, post-fixed overnight in 4% PFA, and subsequently cryoprotected in graded sucrose solutions (10%, 20%, and 30%). For gross visualization of hematoma formation and resolution, brains were sectioned into serial 1-mm-thick coronal slices using a mouse brain matrix. For histological analyses, tissues were embedded, frozen, and sectioned into 40-μm-thick sections using a cryostat. Sections were either processed immediately for staining or stored at −20 °C in cryoprotectant solution consisting of 30% ethylene glycol and 30% glycerol in PBS.

### Magnetic resonance imaging (MRI) and lesion volume analysis

Longitudinal magnetic resonance imaging (MRI) was performed on day 3 of Sham and ICH induction. Imaging was carried out using a 7T small-animal MRI scanner (MR Solutions, Guildford, UK) equipped with a mouse brain coil. During MRI acquisition, mice were anesthetized with 1.5–2% isoflurane in medical-grade oxygen, and body temperature was maintained at 37 °C using a circulating warm-water heating system. Respiratory rate and physiological status were monitored throughout the imaging procedure to ensure stable anesthesia and minimize variability that could influence hemorrhage progression.

T2-weighted fast spin-echo (T2w FSE) images were acquired to visualize hematoma formation, peri-lesional edema, and surrounding tissue damage. Typical imaging parameters included repetition time (TR) =5000ms, echo time (TE)=45 ms, slice thickness=1.0 mm, field of view=220X220 mm^2^, and matrix size= 256X238.

For quantitative analysis, lesion volume was defined as the hypointense region on T2-weighted images encompassing the hematoma core and associated peri-lesional edema within the ipsilateral hemisphere. Lesion volumes were quantified from serial axial slices using MicroDicom software. Regions of interest (ROIs) were manually delineated on each slice by an investigator blinded to experimental groups, and the total lesion volume was calculated by summing the lesion area across slices and multiplying by slice thickness. Hematoma resolution and treatment effects were evaluated by comparing lesion volumes across time points and experimental groups.

### Immunofluorescence and immunohistochemistry

Brain sections were permeabilized with 0.3% Triton X-100 and blocked with 5% bovine serum albumin before incubation with primary antibodies against neuronal (MAP2 1:250), astrocytic (GFAP 1:500), oligodendrocyte (Olig2 1:500), and microglial (Iba1 1:500, CD68 1:250) markers. DNA damage responses were assessed using antibodies against γH2AX (1:250). For immunofluorescence, sections were incubated with appropriate fluorophore-conjugated secondary antibodies and counterstained with DAPI. Fluorescence and brightfield images were acquired using an Olympus slide scanner and a FluoView FV3000 confocal microscope (Olympus). Confocal images were collected as Z-stacks and processed to generate maximum intensity projections for analysis. Colocalization analyses were performed in peri-hematomal regions^31^.

For immunohistochemistry, selected sections were processed using the ABC method. Following incubation with biotinylated secondary antibodies, signal detection was performed using a horseradish peroxidase–conjugated ABC reagent and visualized with DAB substrate. Brightfield images were captured using an Olympus slide scanner.

### Senescence-associated β-galactosidase (SA-β-Gal) staining

SA-β-Gal staining was performed on floating brain sections according to the manufacturer’s instructions (Senescence β-Galactosidase Staining Kit, 9860). Briefly, sections were incubated with X-gal staining solution at 37 °C overnight. Blue precipitate formation was used to identify senescent cells. To assess cell-type-specific senescence, SA-β-Gal staining was combined with immunohistochemistry for neuronal (NeuN) and oligodendrocyte (Olig2) markers using the ABC method. Quantification of senescent cells was performed in peri-hematomal regions.

### Statistical analysis

Data are presented as mean ± SEM. Statistical analyses were performed using GraphPad Prism. Normality was assessed before analysis. Comparisons among multiple groups were conducted using one-way or two-way ANOVA followed by appropriate post hoc tests. Statistical significance was defined as p < 0.05.A p > 0.05 was considered non-significant (ns).

## Results

### Robust induction of ICH triggers marked astroglial and microglial activation

To establish the experimental ICH model, autologous blood was stereotaxically injected into the striatal region. A schematic representation of the injection paradigm is shown in Figure 1a. Gross coronal brain sections confirmed successful hematoma formation in the ipsilateral hemisphere of ICH animals (Figure 1b). The hematoma was consistently localized to the targeted basal ganglia region, validating the reproducibility of the model (Figure 1b). Immunofluorescence analysis further characterized glial responses following ICH. DAPI/GFAP staining revealed minimal astrocytic reactivity in sham brains, with GFAP primarily confined to resting astrocytes exhibiting fine processes. In contrast, ICH brains demonstrated robust GFAP upregulation in the peri-hematomal region, characterized by hypertrophic astrocytes with dense and thickened processes, indicative of reactive astrogliosis (Figure 1c). Similarly, DAPI/Iba1 staining showed sparsely and evenly distributed microglia in sham animals, consistent with a resting state. Following ICH, Iba1 immunoreactivity was markedly increased in the ipsilateral hemisphere, with microglia displaying enlarged cell bodies and retracted processes, consistent with activation (Figure 1c). The accumulation of Iba1-positive cells surrounding the hematoma further indicates a strong inflammatory response. Collectively, these findings confirm successful induction of ICH and demonstrate pronounced astroglial and microglial activation in the peri-lesional region following hemorrhagic injury.

**Figure 1.**
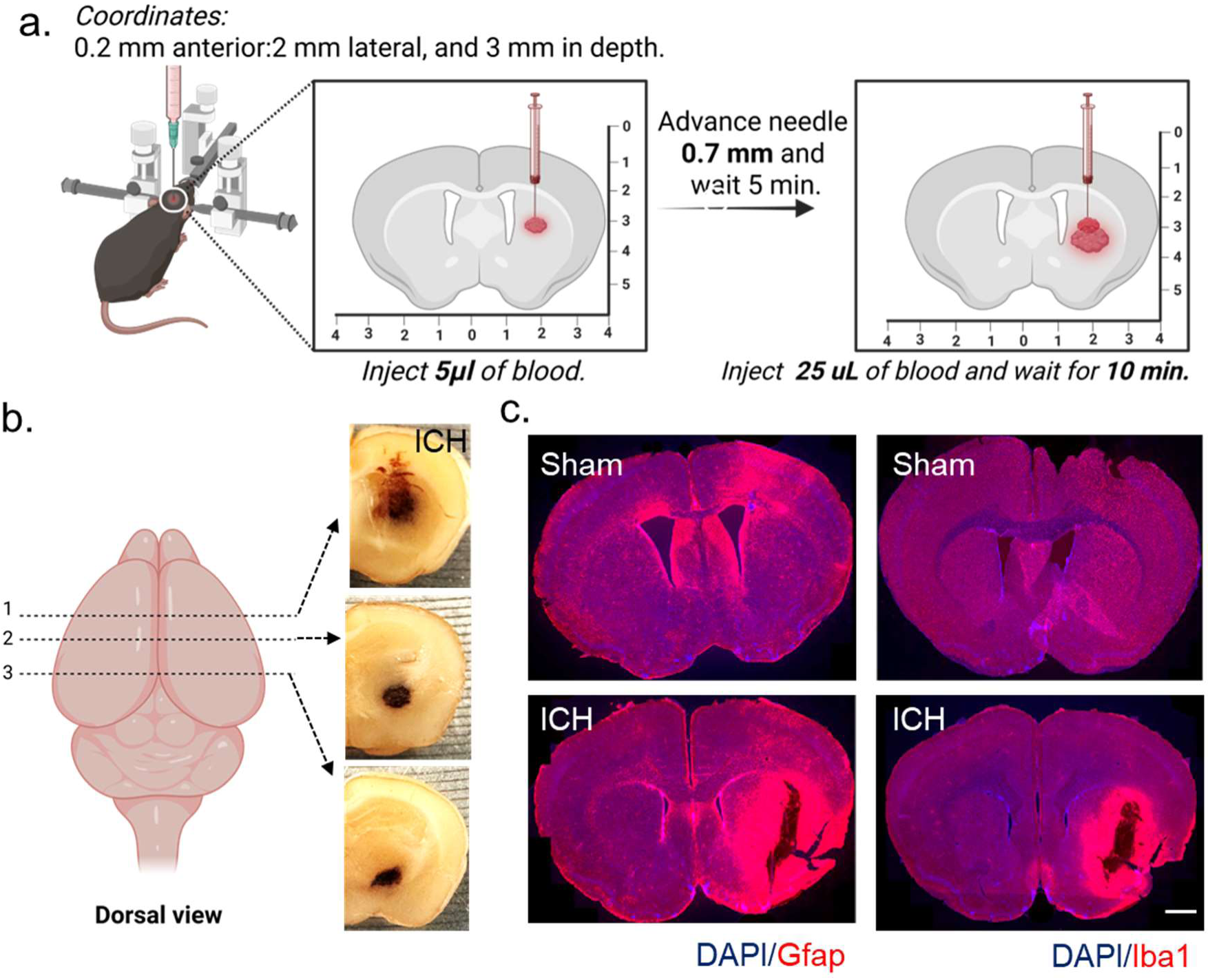
Experimental induction of intracerebral hemorrhage (ICH) in mice and associated astroglial and microglial activation. (a) Schematic illustration of the stereotaxic autologous blood injection model used to induce ICH in the mouse basal ganglia. Autologous whole blood (30 μL), collected from the tail artery, was stereotaxically injected into the right striatum at coordinates relative to bregma (AP: 0.2 mm anterior; ML: 2.0 mm lateral; DV: 3.0 mm depth). Blood was delivered via a Hamilton microsyringe at 2 μL/min using the double-injection model. Following injection, the needle was left in place for 10 min to prevent reflux before slow withdrawal. (b) Representative gross coronal brain sections from sham and ICH mice showing hematoma formation following autologous blood injection. Brains were harvested and sectioned into 1-mm-thick coronal slices using a mouse brain matrix to visualize the location and extent of the hemorrhagic lesion in the ipsilateral hemisphere. The accompanying schematic illustrates the anatomical orientation of coronal sectioning across the mouse brain, indicating the approximate rostro-caudal levels used for histological analysis. (c) Representative immunofluorescence images of coronal brain sections from sham and ICH groups showing astroglial and microglial activation in the peri-hematomal region. Sections were stained with DAPI (blue) to label nuclei and either GFAP (red) to detect astrocytes or Iba1 (red) to detect microglia. Images were acquired using fluorescence microscopy. Increased GFAP and Iba1 immunoreactivity in the peri-lesional region of ICH brains indicates robust activation of astrocytes and microglia compared with sham controls. Scale bar = 1 mm.

### ICH induces cell–type–specific DNA damage and senescence in neurons and oligodendrocytes

To determine whether ICH induces DNA damage in specific neural cell populations, brain sections were co-stained for γH2AX together with Olig2 or MAP2 (Figure 2a & b). In sham brains, γH2AX immunoreactivity was minimal and largely absent in Olig2-positive oligodendrocytes and MAP2-positive neurons (Figure 2a & b: left side panel). In contrast, ICH brains exhibited a marked increase in γH2AX-positive nuclei within the peri-hematomal region. Overlay images demonstrated clear colocalization of γH2AX with both Olig2- and MAP2-positive cells, indicating that hemorrhagic injury induces robust DNA double-strand breaks in oligodendrocytes and neurons (Figure 2a & b: right side panel). Quantitative analysis (violin plots) confirmed a significant increase in γH2AX-positive cells following ICH compared to sham controls.

**Figure 2.**
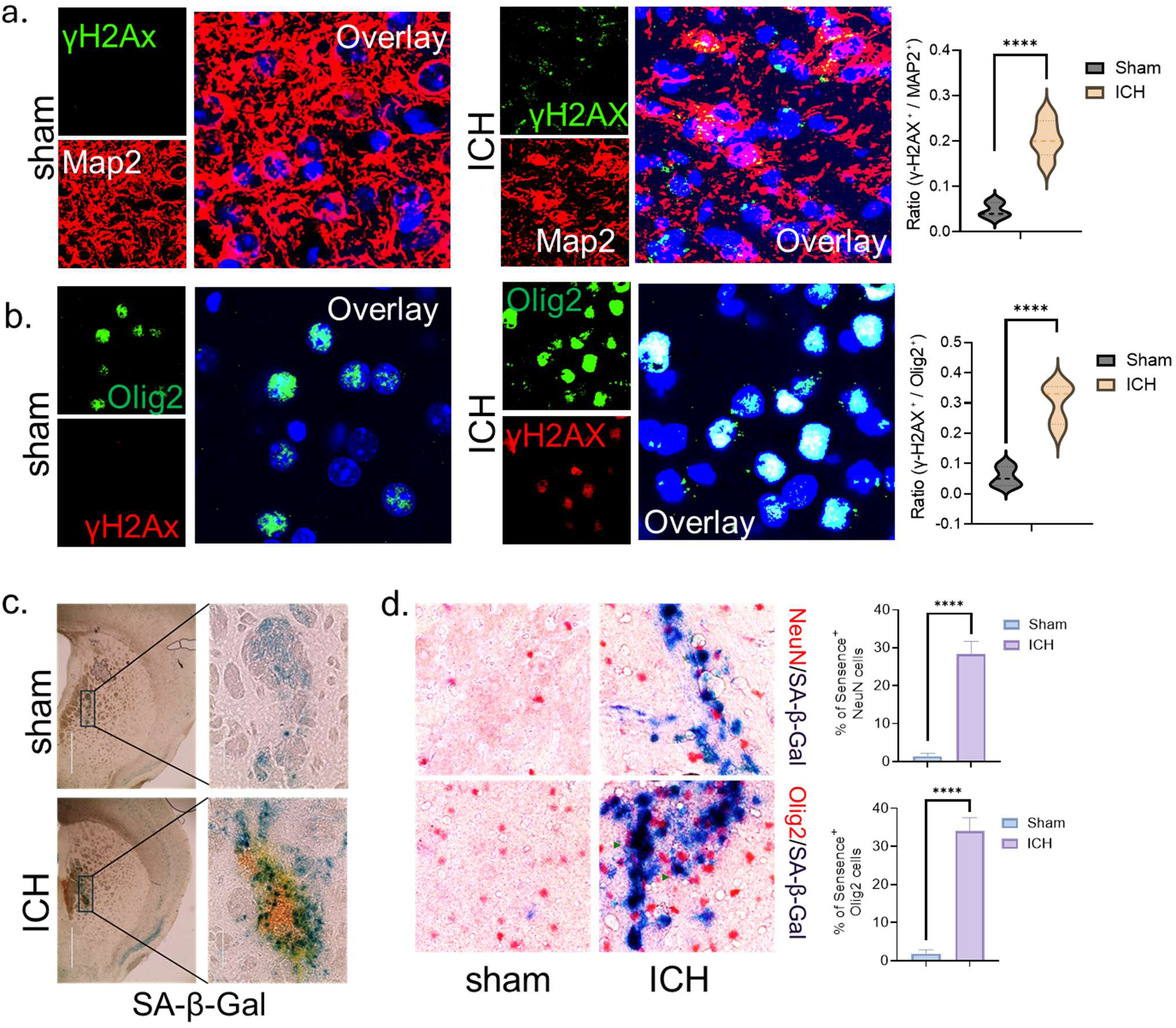
Peri-hematomal neurons and oligodendrocytes exhibit elevated DNA double-strand breaks and senescence following ICH. (a, b) Representative confocal images of coronal brain sections from sham and ICH mice collected at 3 days post-ICH showing γH2AX immunostaining as a marker of DNA double-strand breaks in MAP2-positive neurons and Olig2-positive oligodendrocytes (DAPI, blue). Overlay images demonstrate clear colocalization of γH2Ax-positive nuclei with both MAP2-positive neurons and Olig2-positive oligodendrocytes within the peri-hematomal region adjacent to the hematoma core. Cryosections (40 μm thickness) were stained with antibodies against γH2AX together with MAP2 or Olig2. Images were acquired using a confocal microscope. Scale bar = 20 μm. Violin plots show quantification of the ratio of γH2AX-positive/MAP2-positive cells and γH2AX-positive/Olig2-positive cells. (c) Senescence-associated β-galactosidase (SA) staining demonstrates increased accumulation of senescent cells (blue) in the ipsilateral (injured) hemisphere compared with sham controls at 3 days post-ICH. The SA-β-Gal staining is seen within and surrounding the hemosiderin deposits (brown). Staining was performed on floating brain sections according to the manufacturer’s protocol. Representative images are shown. Scale bar = 100 μm. (d) Co-staining with NeuN (neuronal marker) or Olig2 (oligodendrocyte marker) confirms the presence of senescent neurons and oligodendrocytes in the peri-hematomal region. Quantification shows a significant increase in SA-β-Gal–positive neuronal and oligodendrocyte populations following ICH compared with sham controls. Colocalization was assessed by counting double-positive cells within the peri-lesional region. Quantification was performed from six animals per group (n = 6) within defined peri-lesional regions surrounding the hematoma. Data represent mean values obtained from independent animals in each experimental group. Statistical analysis was performed using one-way ANOVA. Error bars indicate mean ± SEM. Significance was defined as follows: ns, P > 0.05; *P < 0.05; **P < 0.01; ***P < 0.001; ****P < 0.0001.

Given that persistent DNA damage can promote cellular senescence, we next assessed SA-β-Gal activity. Low-magnification images identified the peri-lesional region for analysis (Figure 2c), and higher magnification revealed a substantial accumulation of SA-β-Gal–positive cells in ICH brains compared with sham animals (Figure 2d). Co-staining with NeuN or Olig2 demonstrated that SA-β-Gal activity was present in both neurons and oligodendrocytes (Figure 2d). Quantification showed a significant increase in the proportion of NeuN- and Olig2-positive cells that were SA-β-Gal positive after ICH (Figure 2d). Together, these findings demonstrate that ICH triggers pronounced DNA damage and promotes senescence-like phenotypes in both neuronal and oligodendroglial populations within the injured hemisphere (Figure 2).

### ICH induces HO-1 expression predominantly in neurons

To identify the cellular source of HO-1 induction following ICH, brain sections were co-stained for HO-1 together with MAP2, GFAP, or Iba1, with DAPI used for nuclear labeling. In sham brains, HO-1 immunoreactivity was minimal across all cell types (Supplementary Figure 1a, b & c). Following ICH, a marked increase in HO-1 expression was observed in the peri-hematomal region (Supplementary Figure 1a, b & c). Overlay images demonstrated prominent colocalization of HO-1 with MAP2-positive neurons, indicating that neuronal populations represent the primary source of HO-1 induction after hemorrhagic injury (Supplementary Figure 1a). In contrast, only minimal colocalization of HO-1 was observed with GFAP-positive astrocytes (Supplementary Figure 1b) and Iba1-positive microglia (Supplementary Figure 1c), suggesting limited HO-1 induction in these glial populations at the examined time point. Collectively, these findings indicate that HO-1 expression following ICH is predominantly localized to neurons, with comparatively minor contribution from astrocytes and microglia, highlighting a neuron-specific stress response in the injured brain.

### DEF-OAC-PEG reduces hematoma burden following ICH

To evaluate the therapeutic efficacy of DEF-OAC-PEG in ICH, mice (n=6) underwent stereotaxic autologous blood injection on day 0, followed by systemic administration of DEF-OAC-PEG (2 mg/kg, intraperitoneal). The first dose was administered 2 h after surgery, followed by dosing on alternate days from day 0 to day 6. Animals were sacrificed at either day 3 or day 7 for endpoint analyses. The overall experimental timeline is illustrated schematically in Figure 3a.

**Figure 3.**
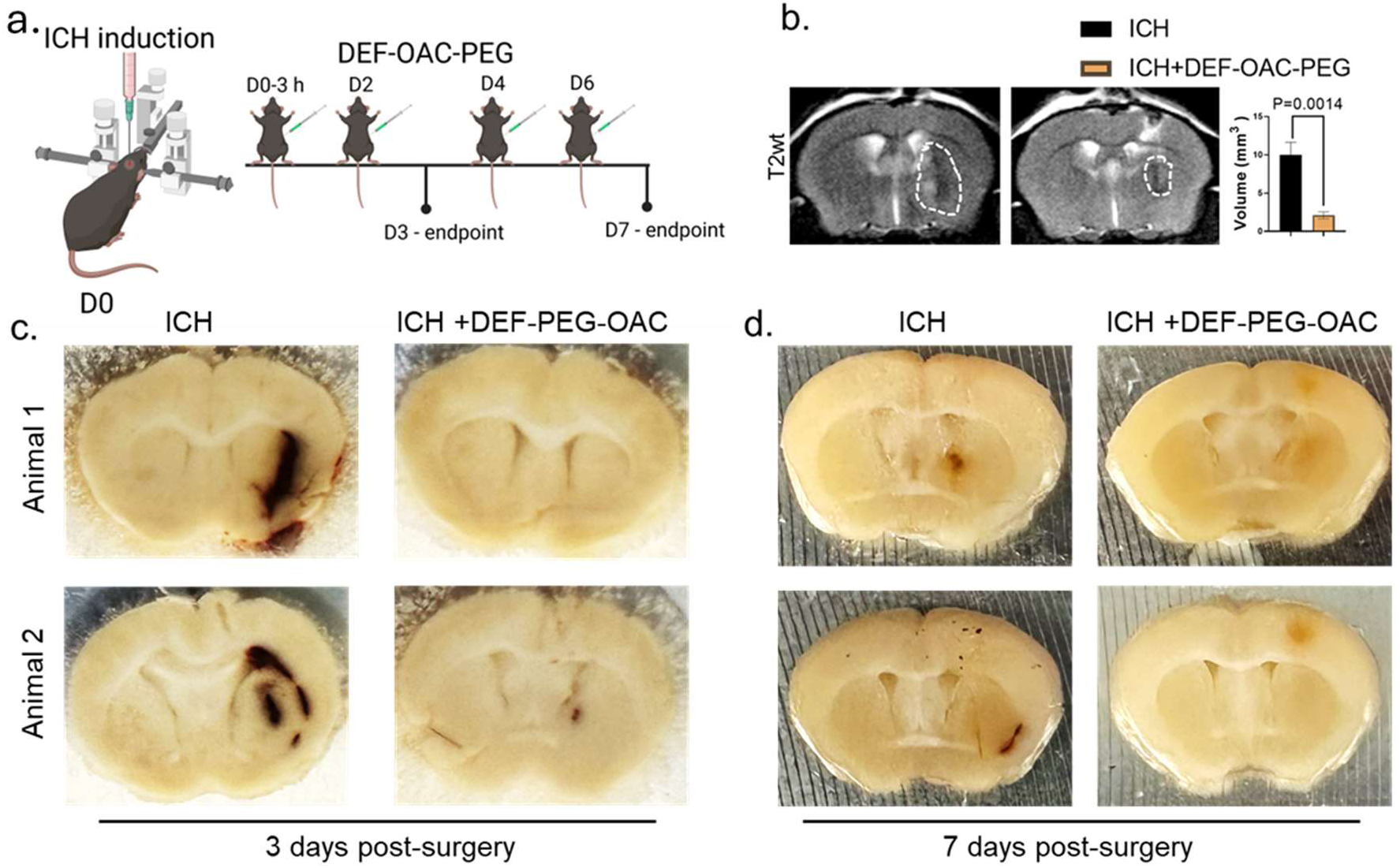
DEF-OAC-PEG treatment accelerates hematoma resolution and reduces lesion volume following ICH. (a) Schematic timeline illustrating ICH induction on day 0 followed by systemic administration of DEF-OAC-PEG nanozymes (2 mg/kg, intraperitoneal) at indicated time points from day 0 (D0) to day 6 (D6). Experimental endpoints were collected at day 3 (D3) and day 7 (D7). (b) Representative T2-weighted MRI images from ICH and ICH + DEF-OAC-PEG groups at D3 post-ICH show the lesion area (dotted outline) in the ipsilateral hemisphere. MRI imaging was performed using a small-animal MRI system, and lesion volumes were quantified from serial axial images using MicroDicom software by an investigator blind to the experimental groups. (c, d) Representative gross coronal brain sections from ICH and ICH + DEF-OAC-PEG groups collected at D3 (left side) and D7 (right side) demonstrate prominent hematoma formation in untreated ICH brains and reduced hemorrhagic area in DEF-OAC-PEG–treated animals. Brains were sectioned into 1 mm coronal slices using a brain matrix to visualize hematoma localization and resolution across time points. Data represent mean values obtained from independent animals in each experimental group. Statistical analysis was performed using one-way ANOVA. Bars represent mean ± SEM. Significance was defined ****P < 0.0001.

MRI analysis demonstrated a well-defined hypointense lesion in the ipsilateral hemisphere of untreated ICH animals (Figure 3b), corresponding to the hematoma region. In contrast, DEF-OAC-PEG–treated mice exhibited a visibly reduced lesion volume at the day 3 endpoint, as indicated by the diminished area within the outlined region (Figure 3b). Gross coronal brain sections further confirmed these findings. Untreated ICH brains displayed prominent hemorrhagic lesions localized to the striatum, characterized by extensive blood accumulation (Figure 3c). In comparison, brains from ICH + DEF-OAC-PEG–treated animals showed markedly reduced hematoma size and improved tissue integrity (Figure 3c) at day 3. These observations were consistent across multiple sections. By day 7, untreated ICH brains exhibited gradual blood clearance, consistent with the natural resolution phase of hemorrhage, accompanied by increased hemosiderin deposition indicative of hemoglobin breakdown and iron accumulation (Figure 3d).Importantly, DEF-OAC-PEG treatment demonstrated persistent reduced residual blood accumulation compared with untreated ICH at both days 3 and 7. Collectively, these results demonstrate that DEF-OAC-PEG treatment attenuates hematoma burden.

### DEF-OAC-PEG localizes to mitochondria and is internalized by CD68-positive phagocytic cells following ICH

To determine the cellular uptake and distribution of DEF-OAC-PEG in the injured brain, mice received systemic (intraperitoneal) administration following ICH induction. Brain sections were co-stained with anti-PEG to detect the nanozyme (targeting covalently bonded PEG to the nanoparticle), TOM20 as a mitochondrial outer membrane marker, and DAPI for nuclear labeling. TOM20 was selected based on our prior in vitro observations suggesting preferential mitochondrial localization of the nanozyme^28^.

Immunofluorescence imaging revealed ananti-PEG signal in uninjured controls (Figure 4a, first row) consistent with uptake of the DEF-OAC-PEG in normal brain. In the ICH-hemisphere, there was robust anti-PEG signal within the peri-lesional region. Overlay images demonstrated clear colocalization of anti-PEG with TOM20, indicating mitochondrial association of the nanozyme in brain cells following ICH (Figure 4b, second row). Notably, an anti-PEG signal was detected in both ipsilateral and contralateral cortical regions, and it also colocalized with TOM20, suggesting widespread mitochondrial engagement beyond the immediate peri-hematomal area ICH (Figure 4b).

**Figure 4.**
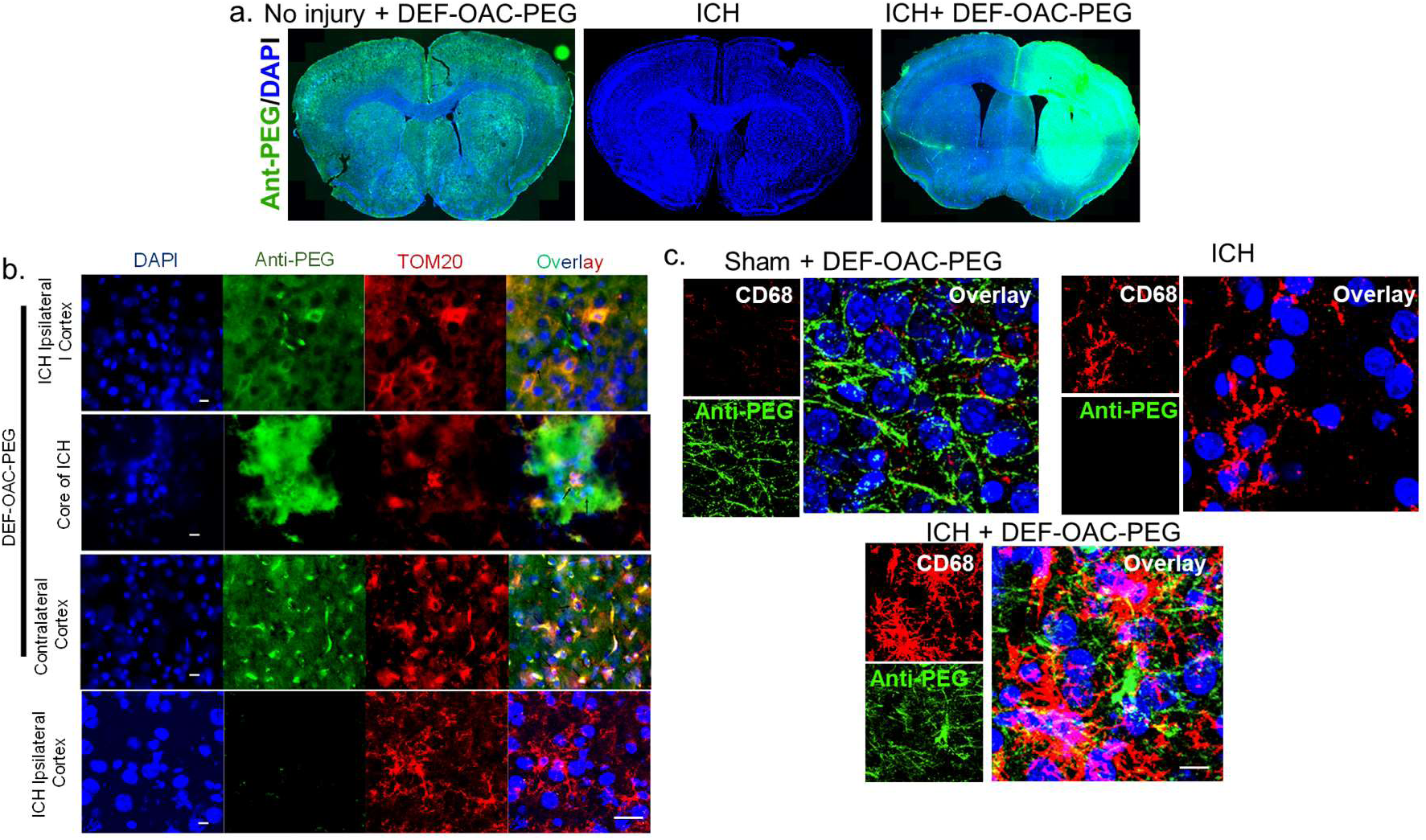
Brain distribution and cellular uptake of DEF-OAC-PEG nanozymes with mitochondrial localization and association with CD68-positive phagocytes after ICH. (a) Representative fluorescence images of coronal brain sections showing the distribution of DEF-OAC-PEG nanozymes following systemic administration. The left image shows a section from a normal mouse injected with DEF-OAC-PEG, demonstrating cellular uptake of the nanozyme within brain tissue. The middle image shows an untreated ICH brain, where minimal anti-PEG signal is observed. The right image shows a section from an ICH mouse treated with DEF-OAC-PEG, where increased accumulation of anti-PEG signal is observed around the peri-hematomal region, indicating preferential localization of nanozymes near the injury site. Sections were stained with DAPI (blue) and anti-PEG (green) to detect the nanozyme. (b) Representative confocal images of coronal brain sections from ICH + DEF-OAC-PEG–treated mice collected at 3 days post-ICH stained with DAPI (blue), anti-PEG (green) to detect the nanozyme, and TOM20 (red) as a mitochondrial marker. Overlay images demonstrate intracellular localization of DEF-OAC-PEG within peri-lesional brain cells and partial colocalization with TOM20-positive mitochondria, indicating mitochondrial association of the nanozyme following systemic administration. Scale bar = 50 μm. (c) Co-immunostaining with CD68 (red), a marker of phagocytic microglia/macrophages, and anti-PEG (green) demonstrates uptake of DEF-OAC-PEG nanozymes within CD68-positive phagocytic cells in the peri-hematomal region. Overlay images show intracellular PEG signal within CD68-positive cells, indicating active internalization of the nanozyme by phagocytic cells after ICH. Scale bar = 50 μm.

To explore whether phagocytic cells contribute to nanozyme distribution and hematoma resolution, we assessed CD68 expression, a marker of phagocytic microglia/macrophages^32^. ICH induced marked activation of CD68-positive cells, which was further evident in DEF-OAC-PEG–treated animals. Co-immunostaining for CD68 and anti-PEG demonstrated intracellular localization of the nanozyme within CD68-positive phagocytic cells, indicating active internalization in the injured hemisphere ICH (Figure 4c).

Collectively, these findings demonstrate that DEF-OAC-PEG accumulates in the peri-hematomal region, associates with mitochondria, and is internalized by CD68-positive phagocytic cells, supporting its engagement within metabolically active and inflammatory compartments following ICH.

### DEF-OAC-PEG reduces DNA damage in oligodendrocytes and neurons following ICH

To evaluate whether DEF-OAC-PEG mitigates ICH-induced DNA damage in specific neural cell populations, brain sections were co-stained for γH2AX together with Olig2 or MAP2. In sham brains, γH2AX immunoreactivity was minimal in both Olig2- and MAP2-positive cells (Figure 5a & b). ICH markedly increased γH2AX-positive nuclear foci within the peri-hematomal region, with clear colocalization observed in both oligodendrocytes and neurons (Figure 5a & b).

**Figure 5.**
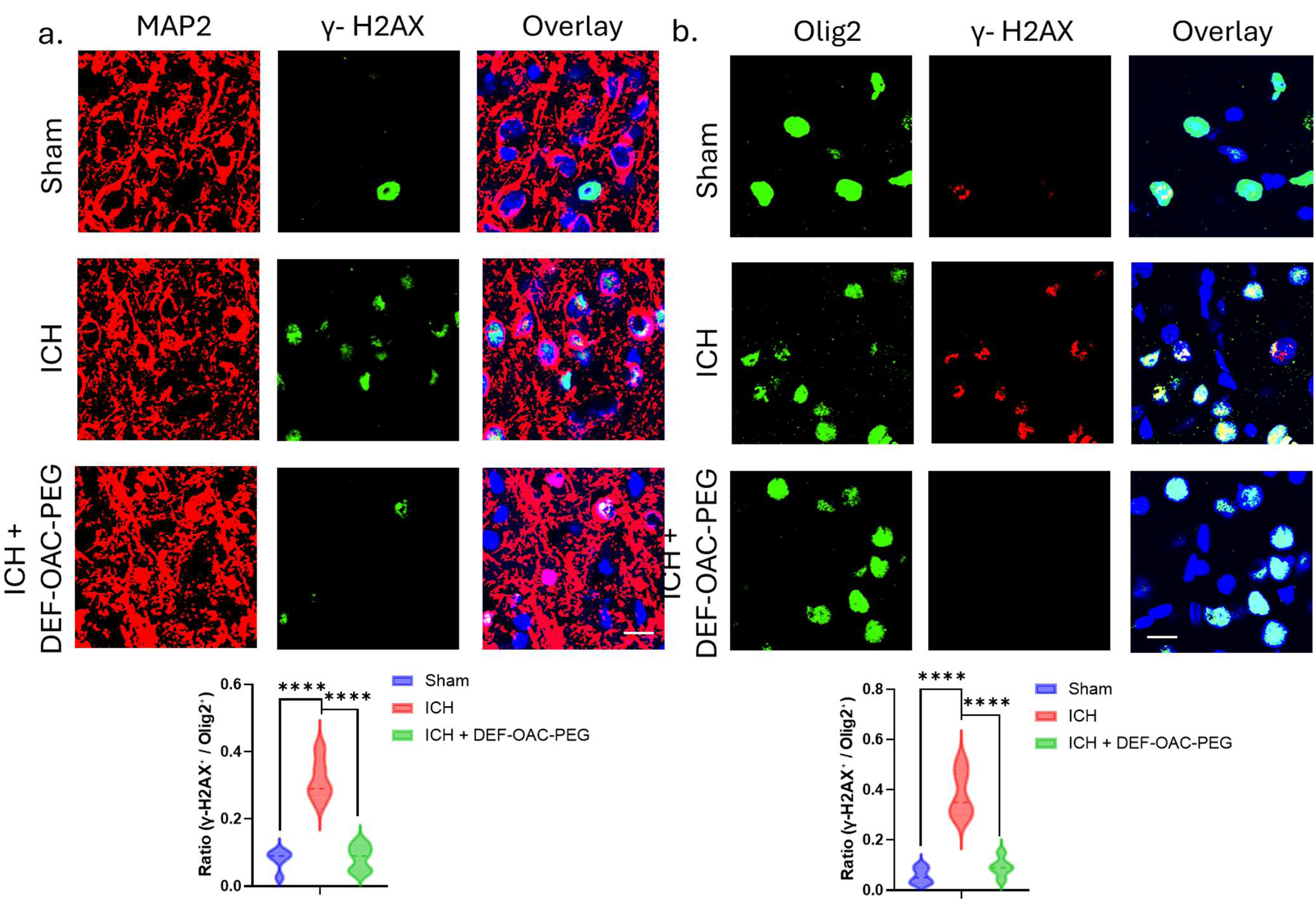
DEF-OAC-PEG treatment mitigates ICH-induced DNA damage in neurons and oligodendrocytes. (a) Representative confocal images of coronal brain sections from Sham, ICH, and ICH + DEF-OAC-PEG groups collected at 3 days post-ICH stained for Olig2 (green; oligodendrocyte lineage marker), γH2AX (red; marker of DNA double-strand breaks), and DAPI (blue; nuclei). Images show increased γH2AX-positive nuclei in Olig2-positive oligodendrocytes within the peri-hematomal region following ICH, which is markedly reduced in DEF-OAC-PEG–treated animals. Sections were imaged using confocal microscopy. Scale bar = 50 μm. (b) Representative images of adjacent sections stained for MAP2 (red; neuronal marker), γH2AX (green), and DAPI (blue) in the same experimental groups. ICH markedly increased the number of γH2AX-positive nuclei in MAP2-positive neurons in the peri-hematomal region, whereas DEF-OAC-PEG treatment significantly reduced γH2AX accumulation in these neuronal populations. Violin plots show quantitative analysis of DNA damage expressed as the percentage of γH2AX-positive nuclei within Olig2-positive oligodendrocytes or MAP2-positive neurons. Quantification was performed from six animals per group (n=6) within defined peri-lesional regions. Data represent mean values obtained from independent animals in each experimental group. Statistical analysis was performed using one-way ANOVA. Error bars indicate mean ± SEM. Significance was defined as ****P< 0.0001.

Treatment with DEF-OAC-PEG substantially reduced γH2AX signal intensity and the number of γH2AX-positive nuclei in both cell populations compared with untreated ICH animals. Overlay images demonstrated decreased colocalization of γH2AX within Olig2- and MAP2-positive cells following treatment (Figure 5a & b).

Quantitative analysis (violin plots) of the ratio of γH2AX-positive/Olig2-positive and γH2AX-positive/MAP2-positive cells confirmed a significant increase in DNA damage after ICH, which was significantly attenuated by DEF-OAC-PEG administration (Figure 5a & b).

These findings indicate that DEF-OAC-PEG effectively reduces ICH-induced DNA damage in both oligodendrocytes and neurons, supporting its neuroprotective potential in the injured brain.

### DEF-OAC-PEG treatment attenuates ICH-induced cellular senescence in the peri-hematomal region

To determine whether ICH induces cellular senescence and whether DEF-OAC-PEG treatment mitigates this response, SA-β-Gal staining was performed on brain sections from sham, ICH, and ICH + DEF-OAC-PEG groups. Minimal SA-β-Gal–positive staining was detected in sham brain sections, whereas ICH resulted in a pronounced accumulation of senescent cells within the peri-hematomal region. Quantitative analysis confirmed a significant increase in SA-β-Gal–positive cells following ICH compared with sham controls. Importantly, treatment with DEF-OAC-PEG markedly reduced the number of SA-β-Gal–positive cells relative to untreated ICH animals, indicating that nanozyme administration significantly attenuates the senescence response associated with hemorrhagic brain injury (Figure 6).

**Figure 6.**
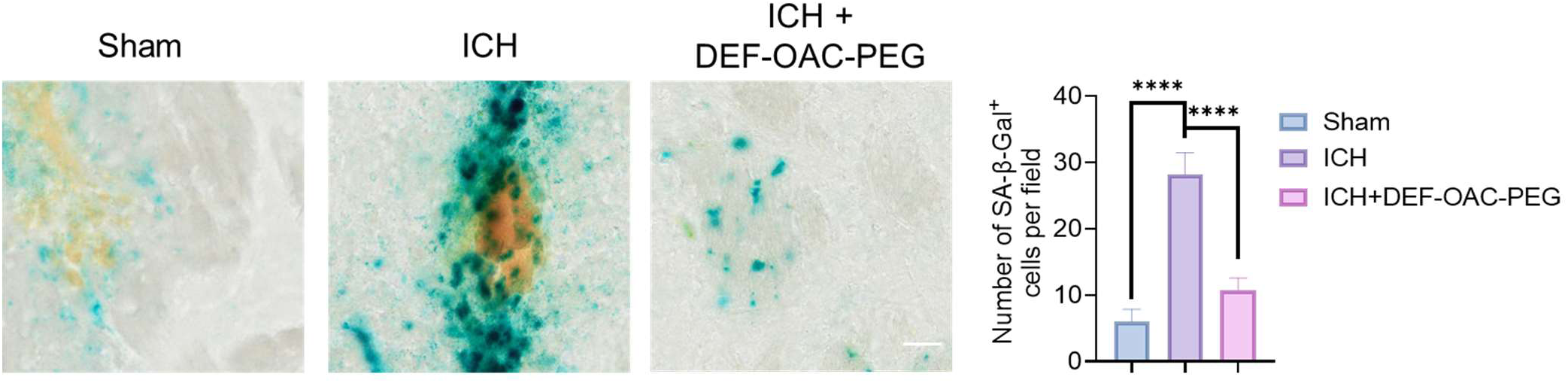
DEF-OAC-PEG treatment reduces senescent cell accumulation in the peri-hematomal region following ICH. Representative images of SA-β-Gal staining in brain sections from Sham, ICH, and ICH + DEF-OAC-PEG groups. Blue precipitate indicates SA-β-Gal–positive senescent cells. Minimal SA-β-Gal staining was observed in sham brain sections, whereas robust senescent cell accumulation was detected in the peri-hematomal region following ICH. Treatment with DEF-OAC-PEG markedly reduced the number of SA-β-Gal–positive cells compared with untreated ICH animals. Quantification of SA-β-Gal–positive cells in the peri-lesional region demonstrates a significant increase in cellular senescence following ICH, which is attenuated by DEF-OAC-PEG treatment. Scale bar = 50 μm. Data represent mean values obtained from independent animals in each experimental group. Statistical analysis was performed using one-way ANOVA. Error bars indicate mean ± SEM. Significance was defined as follows: Significance was defined as ****P < 0.0001.

## Discussion

In this study, we demonstrate that experimental ICH induced by autologous whole-blood injection triggers robust glial activation, neuronal HO-1 induction, DNA damage, and senescence-like changes in vulnerable neural cell populations, particularly neurons and oligodendrocytes. Importantly, systemic treatment with the multifunctional iron-chelator/antioxidant nanozyme DEF-OAC-PEG reduced hematoma burden, promoted hematoma clearance, localized to mitochondria, accumulated in peri-hematomal regions, attenuated DNA damage in neurons and oligodendrocytes, and reduced senescence cell accumulation. Together, these findings provide in vivo evidence that hemorrhagic brain injury is tightly linked not only to classical inflammatory and oxidative pathways, but also to genome instability and senescence-associated pathology, and that a multifunctional catalytic nanozyme approach can beneficially modulate these interconnected mechanisms.

The nanozyme platform used in this study was originally conceptualized by the Kent laboratory to address the complex, multifactorial injury mechanisms that drive hemorrhagic and neurodegenerative brain damage^20,21,27,28,33,34^. The base material, PEG-OAC, was designed as a biocompatible carbon nanoscaffold with intrinsic redox-active properties, enabling broad antioxidant and cytoprotective activity^20,28^. Building on this platform, the nanozyme was further functionalized with the clinically used iron chelator DEF to generate DEF-OAC-PEG, thereby integrating iron sequestration with the intrinsic antioxidant and mitochondrial-protective features of the nanoparticle^20,27^. Initial functional characterization demonstrated that this nanozyme exhibits favorable cellular uptake, brain penetrance, and subcellular localization, including mitochondrial association, while also reducing oxidative injury, preserving metabolic function, and mitigating hemin- and iron-induced injury in cultured neural and vascular cells^20,21,28^. Importantly, prior studies showed that the platform can suppress hemin-induced DNA damage, reduce cellular senescence, and protect against iron-mediated ferroptotic stress, establishing its utility as a multifunctional therapeutic strategy for disorders with oxidative injury, iron overload, mitochondrial dysfunction, and genome instability^20,21^. These foundational studies provided the rationale for advancing DEF-OAC-PEG into the present in vivo investigation of ICH.

A major strength of this study is the use of the two-injection autologous whole-blood injection model, which closely mimics the pathological consequences of blood extravasation into the brain parenchyma and rebleeding, while minimizing confounding inflammatory effects that may arise from non-autologous or exogenous components^29,35,36^. By using the animal’s own blood, this model better isolates the intrinsic toxic effects of hematoma formation, erythrocyte lysis, hemin release, iron accumulation, and endogenous immune activation^37^. This is particularly important in the context of evaluating nanozyme-mediated hematoma resolution and secondary tissue protection, because it reduces the likelihood that the observed effects are driven by artificial inflammatory stimuli unrelated to the core biology of hemorrhagic stroke. The robust astroglial and microglial activation observed here, therefore, likely reflects physiologically relevant responses to blood-derived injury and tissue stress rather than extrinsic immunogenic artifacts (Figure 1c).

Our findings further extend prior in vitro observations by establishing in vivo relevance for the link between hemorrhagic injury, genome damage, and senescence^20,21^. While oxidative injury, lipid peroxidation, and iron-mediated neurotoxicity are well-established features of ICH^38,39^, the contribution of persistent DNA damage and senescence-like phenotypes to secondary brain injury has remained insufficiently explored. The marked increase in γH2AX-positive nuclei in neurons and oligodendrocytes, together with increased SA-β-Gal activity in these populations, supports the idea that ICH induces a broad stress response that includes activation of DNA damage pathways and a senescence-associated phenotype (Figure 2). These findings are consistent with the theory that blood breakdown products, especially hemin and iron, can impair both nuclear and mitochondrial homeostasis and thereby drive long-lasting degenerative consequences beyond the initial mechanical insult^40^. Such processes may be especially relevant to chronic tissue remodeling, white matter damage, and long-term cognitive decline after ICH.

One of the most striking discoveries in this study is that DEF-OAC-PEG treatment was associated with a reduction in hematoma volume and accelerated hematoma clearance (Figure 3). The nanozyme localized within CD68-positive phagocytic microglia/macrophages, suggesting that it may enhance or support endogenous phagocytic mechanisms involved in blood and debris clearance (Figure 4c). This is an intriguing observation because efficient hematoma resolution is a key determinant of tissue recovery, yet the cellular and biochemical processes that govern this response are incompletely understood^8,41,42^. A plausible interpretation is that the nanozyme improves the intracellular redox and iron environment of phagocytic cells, thereby preserving mitochondrial fitness and enhancing their capacity to process heme-rich material. In support of this, our recent studies have revealed the enzymatic role of these nanozymes in improving mitochondrial functions and metabolism^27,28^. However, the precise mechanisms by which DEF-OAC-PEG promotes hematoma clearance remain to be defined.

Accordingly, an important next step will be to identify the mechanistic basis of nanozyme-enhanced blood clearance in greater depth. Future studies should address whether DEF-OAC-PEG alters microglial/macrophage polarization states, lysosomal function, erythrophagocytosis, iron handling pathways, HO-1 signaling, ferritin induction, or mitochondrial bioenergetics within peri-hematomal phagocytes. It will also be important to determine whether enhanced clearance reflects improved survival and function of resident microglia, recruitment of infiltrating macrophages, or both. Such mechanistic studies could reveal new therapeutic entry points for accelerating hematoma resorption while limiting secondary iron toxicity.

Another important implication of this work is that multifunctional nanozyme platform may be particularly well suited for complex disorders such as ICH, where no single pathogenic mechanism predominates^43^. This multifunctionality likely underlies the ability of the DEF-OAC-PEG nanozyme to mitigate both hematoma-associated injury and downstream DNA damage and senescence (Figure 5 & 6). Future optimization of this platform may further improve efficacy and specificity. For example, next-generation nanozymes incorporating alternative therapeutic moieties, including more selective ferroptosis inhibitors, may provide stronger protection against iron-driven lipid peroxidation while potentially reducing off-target effects or toxicity. Such modular redesign could allow better tuning of the nanozyme for different stages of ICH pathology, including acute hematoma toxicity, subacute inflammatory injury, and chronic neurodegenerative conditions.

Although promising, the present study has several limitations. First, the current dataset is focused primarily on histological, imaging, and cellular injury endpoints at relatively early time points, and longitudinal evaluation of senescence and ferroptosis are still pending. These additional studies will be important for fully defining the relationship between nanozyme treatment, iron-dependent cell death pathways, and persistence versus resolution of senescence. Second, while the reduction in hematoma burden and DNA damage is encouraging, the study does not yet establish whether these improvements translate into meaningful functional recovery. Future studies should therefore incorporate comprehensive behavioral and neurological assessments, including motor, sensorimotor, and cognitive outcomes across acute and chronic post-ICH intervals. Third, although the observed mitochondrial localization is consistent with our prior in vitro findings, the subcellular trafficking, retention, metabolism, and clearance of the nanozyme in vivo require further study.

In addition, broader pharmacological and toxicological evaluation will be essential before translation. Dose-response studies, therapeutic window analyses, repeated-dose paradigms, biodistribution profiling, and acute and chronic toxicity assessments in multiple organs will be important next steps. Because ICH patients often present with comorbidities and heterogeneous injury burden, future preclinical testing should also evaluate sex as a biological variable, aging, and delayed treatment paradigms, as well as comparisons with collagenase-based or larger-animal hemorrhage models. These studies will help determine the robustness, safety, and translational feasibility of iron-chelator nanozyme therapy.

In summary, this study provides proof of concept that a multifunctional iron-chelator nanozyme can promote hematoma resolution while simultaneously attenuating secondary genomic damage after ICH. The work highlights the importance of investigating hemorrhagic injury not only through the lens of acute oxidative and inflammatory stress, but also through the interacting frameworks of genome instability, senescence, and ferroptosis-related vulnerability. By linking these processes in an in vivo autologous blood model, our findings provide a strong rationale for further development of nanozyme-based therapeutics aimed at limiting both the acute and delayed neurodegenerative consequences of hemorrhagic stroke.

## Acknowledgements

This research was primarily supported by the National Institute of Neurological Disorders and Stroke (NINDS) of the National Institutes of Health under award number R01NS094535 (T.A.K., J.M.T., and M.L.H.). Additional support for research in the M.L.H., P.J.H., and G.W.B. laboratories was provided by the Walter Foundation Endowment NeuroSpark seed award. M.L.H. also acknowledges support from NIH award RF1NS112719 and from Everett E. and Randee K. Bernal through the Centennial Endowed Chair of the Neurological Institute. T.A.K. is supported by the Welch Foundation (grant BE-0048). The authors thank Pavana Hegde for technical/administrative assistance and Dr. Gillian Hamilton and Dr. Anna Dodson at Houston Methodist Research Institute (Houston, TX) for their help with document editing. Schematic illustrations and graphical abstract images were created using elements adapted from BioRender. The authors also acknowledge the HMRI Microscopy Core and HMRI Radiology/Pre-Clinical Imaging Core for their valuable support and resources used for microscopy and MRI imaging in this study.

## Author Contributions

V.H.M. contributed to the study design, performed the majority of the experiments, conducted data analysis, and co-wrote the manuscript. A.M.M. performed most of the ICH surgeries. M.K. assisted with microscopy, data analysis, and statistical analysis. A.V.L. K.M., and P.J.D. synthesized and provided nanoparticles. S.Z. and Z.L. performed MRI imaging of the animals and assisted with image analysis. E.P. assisted with ICH animal surgeries and optimization of the experimental model. X.T. performed immunohistochemistry. P.J.H. provided critical reagents and contributed important intellectual input. G.W.B. and R.C.R. edited the manuscript and provided valuable insights. T.A.K. and M.L.H. co-supervised the study and contributed to the study design, oversight, data analysis, and interpretation of the results, and finalized the manuscript with important feedback from G.W.B. and P.J.H. All authors discussed the results and contributed to the preparation and revision of the manuscript.

## Data availability statement

Original, raw data from the experiments conducted in this study can be provided upon reasonable request to the corresponding authors.

