## Supplementary material for "Multifunctional nanozyme therapy accelerates hematoma clearance and attenuates genome damage and senescence after intracerebral hemorrhage"

**Supplementary Material accompanying this paper includes one figure.**

**Supplementary Figure. 1**

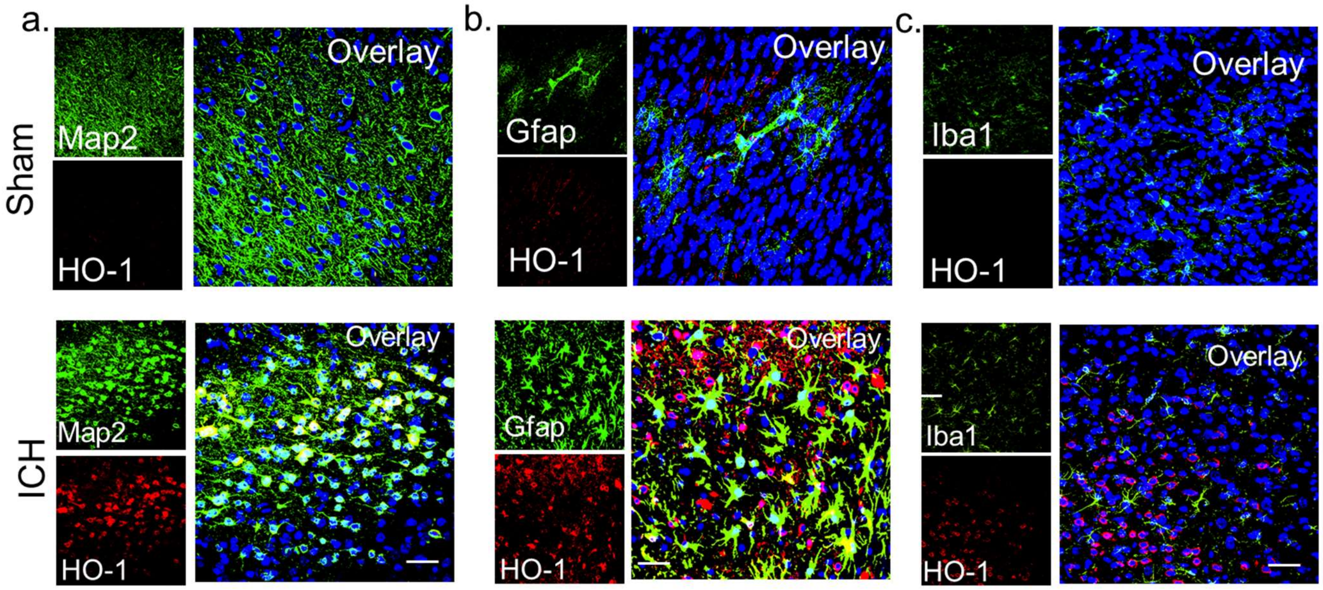


**Supplementary Figure 1. Predominant induction of HO-1 in neurons with minimal expression in astrocytes and microglia following ICH.** Representative confocal images of peri-hematomal brain sections showing cell–type–specific expression of heme oxygenase-1 (HO-1) after intracerebral hemorrhage (ICH). Brain sections were co-stained with HO-1 (red) and cell-type–specific markers, including MAP2 (green; neuronal marker), GFAP (green; astrocyte marker), and Iba1 (green; microglial marker), with DAPI (blue) labeling nuclei. Overlay images demonstrate strong colocalization of HO-1 with MAP2-positive neurons, whereas minimal colocalization is observed with GFAP-positive astrocytes or Iba1-positive microglia. These findings indicate that HO-1 induction following ICH occurs predominantly in neuronal populations within the peri-hematomal region. Scale bar = 100 μm.
